# Enhanced TF binding site maps improve regulatory networks learned from accessible chromatin data

**DOI:** 10.1101/545780

**Authors:** Shubhada R. Kulkarni, D. Marc Jones, Klaas Vandepoele

**Affiliations:** Ghent University, Department of Plant Biotechnology and Bioinformatics, Technologiepark 71, 9052 Ghent, Belgium; VIB Center for Plant Systems Biology, Technologiepark 71, 9052 Ghent, Belgium; Bioinformatics Institute Ghent, Ghent University, Technologiepark 71, 9052 Ghent, Belgium

## Abstract

Determining where transcription factors (TF) bind in genomes provides insights into which transcriptional programs are active across organs, tissue types, and environmental conditions. Recent advances in high-throughput profiling of regulatory DNA have yielded large amounts of information about chromatin accessibility. Interpreting the functional significance of these datasets requires knowledge of which regulators are likely to bind these regions. This can be achieved by using information about TF binding preferences, or motifs, to identify TF binding events that are likely to be functional. Although different approaches exist to map motifs to DNA sequences, a systematic evaluation of these tools in plants is missing. Here we compare four motif mapping tools widely used in the Arabidopsis research community and evaluate their performance using chromatin immunoprecipitation datasets for 40 TFs. Downstream gene regulatory network (GRN) reconstruction was found to be sensitive to the motif mapper used. We further show that the low recall of FIMO, one of the most frequently used motif mapping tools, can be overcome by using an Ensemble approach, which combines results from different mapping tools. Several examples are provided demonstrating how the Ensemble approach extends our view on transcriptional control for TFs active in different biological processes. Finally, a new protocol is presented to effectively derive more complete cell type-specific GRNs through the integrative analysis of open chromatin regions, known binding site information, and expression datasets.

## INTRODUCTION

Plants are exposed to a wide variety of internal and external signals which need to be correctly processed to facilitate growth and development and to trigger defense responses against environmental stimuli. An important mechanism mediating these signal processing pathways is the control of gene expression. Gene expression is regulated by transcription factors (TFs), proteins which often bind to specific, short DNA sequences and influence gene expression. The identification of functional TF binding is an important step in understanding the biological roles of these regulators. Regulatory links between TFs and target genes together form a gene regulatory network (GRN), which can be used to understand the dynamics of plant processes, such as diverse cellular functions, responses to various external stimuli, and organ development (Song et al., 2016; Sparks et al., 2016; Varala et al., 2018).

An early and important step in the characterization of GRNs is understanding TF binding preferences, or motifs, as determining potential binding locations of a TF within a genome assists identification of putative target genes. Advancements in technologies that profile regulatory DNA have successfully characterized the binding preferences of many plant TFs (reviewed in (Franco-Zorrilla and Solano, 2017). Protein binding microarrays, a high throughput experimental technique, determine sequence preferences of TFs by allowing fluorescently labeled proteins to bind to an array of oligonucleotides. Using this technology, TF binding profiles were determined for 63 *Arabidopsis thaliana* (Arabidopsis) TFs, representing 25 TF families, while Weirauch and co-workers identified motifs for >1000 TFs across 131 species (Franco-Zorrilla et al., 2014; Weirauch et al., 2014). Another *in vitro* assay, DNA affinity purification sequencing (DAP-seq), combines *in vitro* expressed TFs with next-generation sequencing of a genomic DNA library. Using this technique, binding profiles for 529 TFs in Arabidopsis have been elucidated (O’Malley et al., 2016). In recent years, numerous TF chromatin immunoprecipitation (ChIP) experiments have been performed, expanding our knowledge of TF binding in plants (Heyndrickx et al., 2014; Song et al., 2016). Collectively, these binding profiles offer an interesting resource to study TF binding in the Arabidopsis genome for over 900 TFs (Kulkarni et al., 2018).

The simplest approach to delineate GRNs from these profiles is by naively mapping the TF motifs to the nearest gene promoter. However, the high rate of false positives when mapping motifs to a DNA sequence, especially if the motif is short and degenerate, results in low specificity to identify functional regulatory events (Baxter et al., 2012). To overcome these issues, additional sources of evidence, such as gene co-regulation or evolutionary sequence conservation, are frequently used to define functional binding sites. Based on the hypothesis that a set of co-regulated genes are regulated by a similar cohort of TFs, identification of over-represented sequences in the promoters of these genes can enrich for functional true positives (Michael et al., 2008; Vandepoele et al., 2009; Hickman et al., 2017; Kulkarni et al., 2018). An alternative approach involves filtering potential binding sites using conservation information over large evolutionary distances. This method assumes that functionally important binding sites will be under purifying selection, and as such, will be conserved between species. Filtering motif matches using this metric significantly reduces the false positive rate (Vandepoele et al., 2006; Haudry et al., 2013; Van de Velde et al., 2014; Burgess et al., 2015; Yu et al., 2015), although it is important to note that not all functional binding events are evolutionary conserved (Muino et al., 2016).

Recent advances in the profiling of open chromatin have increased our understanding of regulatory DNA in Arabidopsis (Zhang et al., 2012; Sullivan et al., 2014; Lu et al., 2017). Combined with cell type-specific nuclear purification, methods such as Assay for Transposase-Accessible Chromatin followed by DNA sequencing (ATAC-Seq) offer unprecedented opportunities to identify cell type-specific TF networks (Lu et al., 2017; Maher et al., 2018; Sijacic et al., 2018). Nevertheless, elucidation of GRNs from chromatin accessibility data requires detailed information about TF binding preferences in order to identify potential binding sites within accessible regions of the genome, and therefore infer TF-target gene regulatory interactions.

Based on the importance of motif mapping to find locations of potentially functional TF binding, in this study we compared four frequently used motif mapping tools and performed a detailed evaluation of their global performance for 40 TFs in Arabidopsis. We evaluated the similarities and differences between these tools at a TF level and found that differences in tool sensitivity and specificity affect the inference of GRNs. By combining the results from two tools into an Ensemble, we were able to improve the identification of TF-target regulatory interactions in different experimental datasets. Using this Ensemble approach we developed a protocol to elucidate cell type-specific GRNs from ATAC-Seq defined accessible genomic regions. The results of this analysis, relative to the original study, offer a more complete view on gene regulation in SAM stem and mesophyll cells in Arabidopsis.

## RESULTS

### Performance of individual motif mapping tools to identify *in vivo* binding events

A wide variety of tools are used in the plant research community to map TF motifs (Supplemental Table S1). We selected and evaluated four frequently used tools to map TF motifs in Arabidopsis: FIMO, Cluster-Buster (CB), matrix-scan (MS), and MOODS (Frith et al., 2003; Turatsinze et al., 2008; Korhonen et al., 2009; Grant et al., 2011). These tools were used to map a set of 66 motifs (corresponding to 40 TFs and 19 TF families; see Supplemental Table S2 for TF motif details) onto the Arabidopsis genome (see “Materials and Methods”). The motifs, mainly derived from protein binding microarrays and DAP-Seq, were selected based on the availability of experimental ChIP-Seq datasets for the profiled TF. The set of TFs included in this analysis have diverse roles in processes such as the cell cycle, flower development, response to light or hormones, and defense responses. Motif matches (referred to as TF binding sites; TFBSs) reported by the different tools were evaluated by counting the number of TFBSs confirmed by ChIP-Seq datasets (precision). Recall for each tool was calculated as the fraction of regions identified by ChIP-Seq that were covered by a motif match, that is, how many target genes are correctly recovered by motif matches (median values of performance statistics in Table 1). FIMO produced the lowest number of motif matches (2.4 million matches versus 19-34 million matches for the other tools) and showed the highest precision among all tools. The median precision for FIMO is 5%, compared to 2.2-2.4% for the other tools (Figure 1A), indicating that FIMO reports a higher fraction of experimentally supported matches. However, recall is low with the FIMO results as a consequence of the tool predicting approximately 10-fold fewer matches relative to the other tools (22% median recall versus 36-48% for the other tools; Figure 1B). Overall, these results suggest that FIMO misses some real TFBSs based on the ChIP-Seq data, considering all matches. Due to the large variation in the total number of matches predicted by each tool, we also evaluated the tool performance considering only the 7000 highest scoring matches (top7000). The size of this subset was chosen to optimize the compromise between precision and recall for CB (see “Materials and Methods”). Using this subset of matches, the median precision and recall of all tools are similar (Figure 1A and 1B). In order to assess the false positive rate for each tool, TF motifs were mapped using shuffled promoter sequences of Arabidopsis genes (see “Materials and Methods”). Due to its stringency, FIMO has the lowest FPR (Figure 1C), while MOODS, which identifies the highest number of matches, has the highest FPR compared to the other tools. Together with the above results, this suggests that many of the matches identified by MOODS are false positives. Overall, the FPR for all tools was below 10%.

**Table 1.**
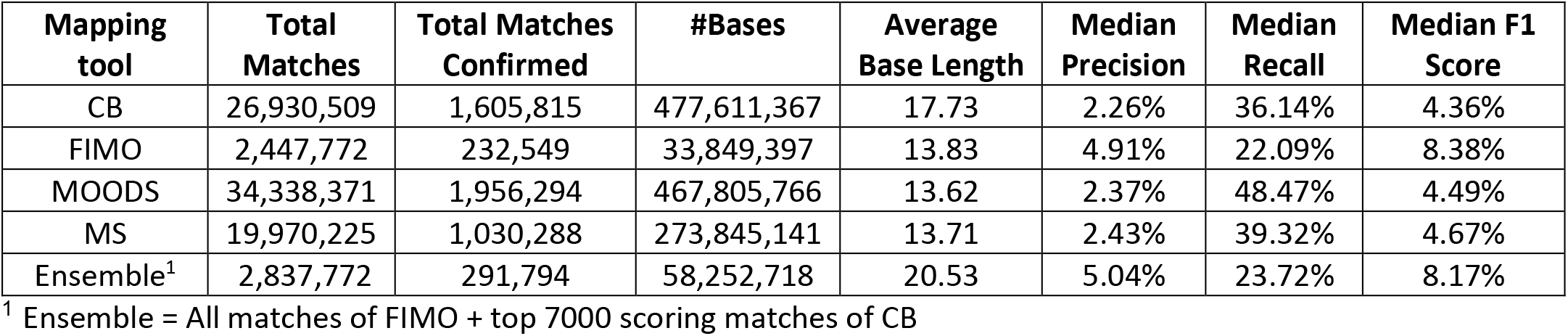
Performance statistics of mapping 66 TF motifs using different motif mapping tools and an Ensemble approach.

**Figure 1.**
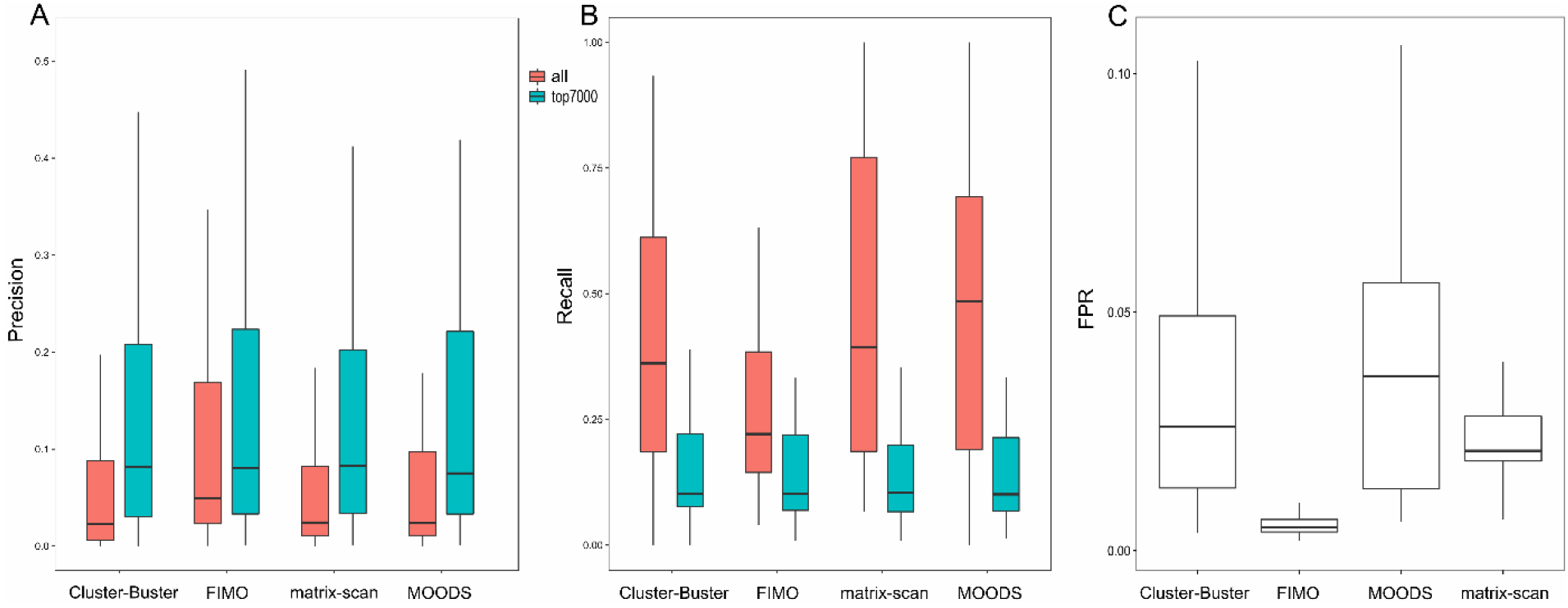
Global performances of mapping tools in Arabidopsis. (A) and (B) Precision and recall of motif matches considering all matches (in red) and top-scoring 7000 matches (in cyan). The boxes indicate the number of matches predicted by each tool. (C) Boxplot showing the false positive rate (FPR) for every tool.

Following the evaluation of mapping tool performances, we next studied the effect of TF motif complexity on the precision and recall values, using the information content (IC) of each motif. Given the clustering pattern in Figure S1, all matches predicted by FIMO and CB were considered for this analysis. To examine the effect of motif complexity on the performance measures, 21 TFs were selected for which more than one motif was available. For CB, for 15 TFs, the F1 score increased with increasing motif complexity (Figure 2). For FIMO, however, this trend was observed for 8 TFs only. FIMO, besides implementing a p-value threshold for calling motif matches, has an internal cut-off to restrict spurious matches when used with low complexity motifs. This additional threshold is likely responsible for the quality of the TFBSs found with FIMO being less dependent upon TF motif complexity. Of the TFs selected for the above analysis, 13 had motifs from different sources, such as CisBP and DAP-Seq. We next checked if the source of motifs had an impact on the performance measures. CisBP motifs, derived from PBMs, were on an average shorter than motifs derived from DAP-Seq (average lengths for CisBP=11.67 and DAP-Seq=14.55) and were less complex (average IC for CisBP=8 and DAP-Seq=10; Supplemental Table S2). For ABI5, AGL15, ERF115, GFB3, HB7 and WRKY33, the F1 score was higher for DAP-Seq motifs compared to CisBP. For the remainder of the TFs, where the complexity between the two motifs did not vary, the F1 scores were similar.

**Figure 2.**
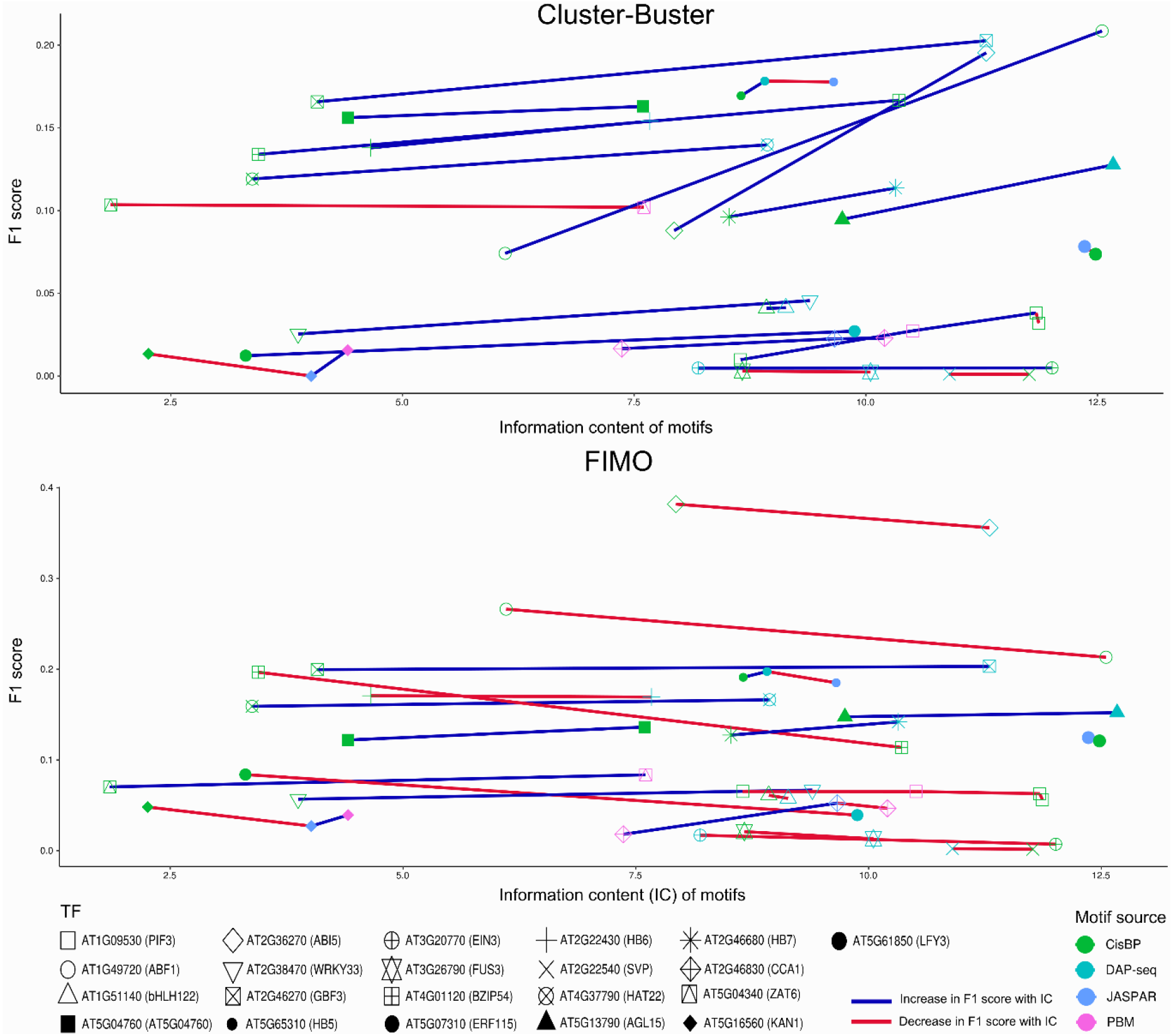
Variation of F1 score with TF motif complexity. Scatterplot showing the effect of motif complexity on F1 scores for CB and FIMO for 21 TFs. Each motif is shaped based on the TF it belongs to and colored based on the source of that motif. Incrase and decrease in F1 score with motif complexity is marked with blue and red lines respectively

### Evaluation of unique motif matches reveals complementary between mapping tools

For the TFs included in our benchmark, the varying recovery of true positive matches suggests that each tool performs differently depending on the complexity of the motif (Figure S1). To investigate the differences between tools further, we compared the motif matches confirmed by ChIP-Seq data for each tool. To account for the large differences in the number of matches reported by each tool, only the top7000 matches per tool and per motif were used in this analysis. Conducting pairwise comparisons between tools reveals that for 12 out of 66 motifs, the TFBSs identified uniquely by CB have high recall rates (Figure S2). This pattern is retained when matches found uniquely by a particular tool, relative to all matches of the other tools combined, are used (Figure S3). For 65 motifs, the recall of the top7000 matches uniquely found with CB was larger than zero, making it the only tool to identify functional matches for 98% of all TF motifs considered in this study (Figure 3A). Moreover, for 21 of 65 motifs, the motif mappings from CB were able to achieve recall values between 10% and 38%, considerably higher than the recall rates of other tools, which did not exceed 10% (Figure 3B). This result highlights that CB is able to identify a unique set of functional matches with high ChIP recovery for 32% of the studied motifs.

**Figure 3.**
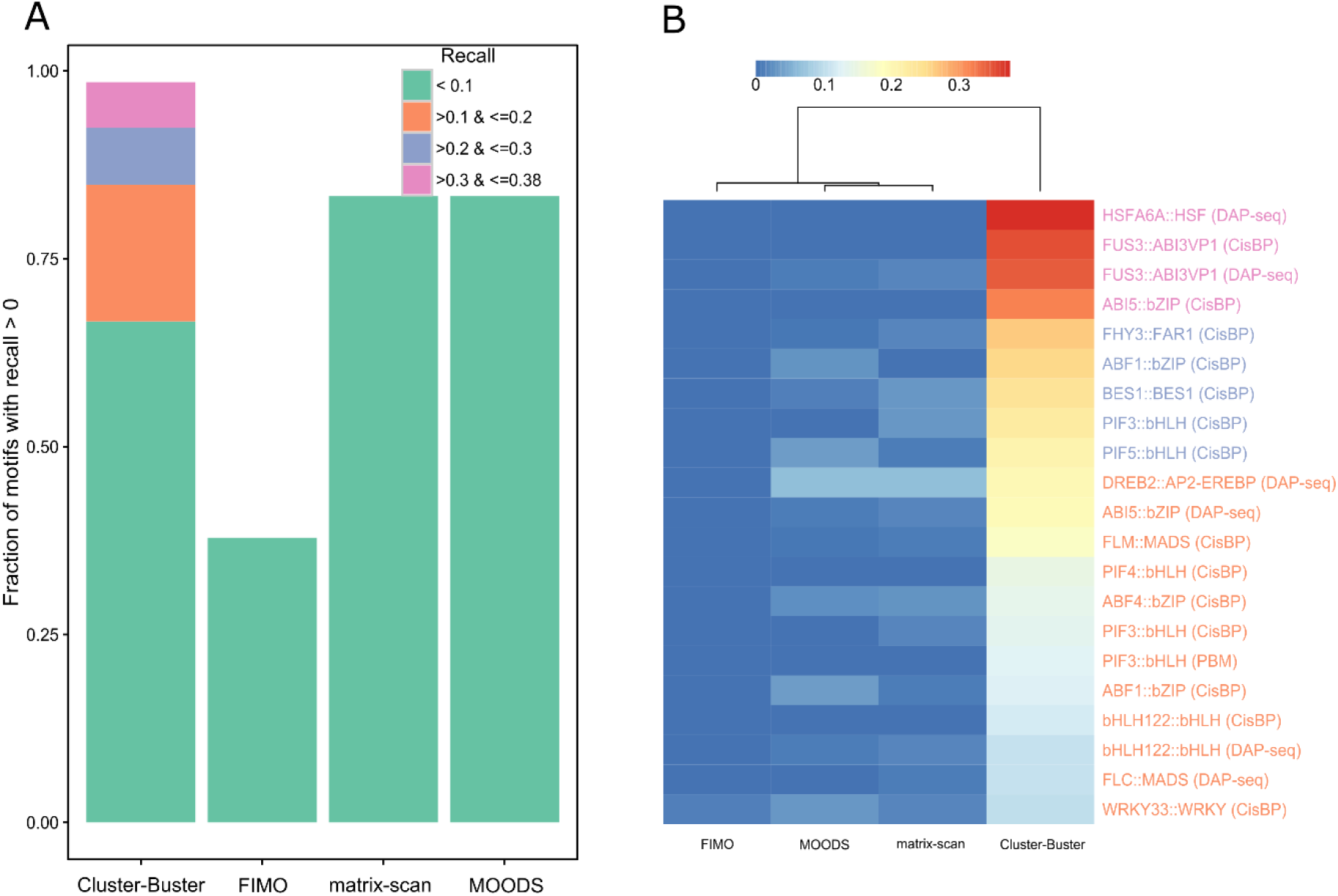
Evaluation of unique matches predicted by each tool. (A) Barplot showing the fraction of TFBS with recall>0 considering unique matches in top7000 of one tool compared to matches of all other tools combined. (B) Heatmap showing the recall for each tool for TFBSs where CB outperforms the other tools (orange, purple and pink series from panel A where recall of CB was >10%).

Another aspect in which the tools differ is in the length of TFBSs reported. The average motif match, considering all matches, is 17.73 bp for CB, whereas for other tools, it is 13.72 bp (Table 1). In some cases, the TFBSs identified by CB are longer than the motifs mapped. This difference is due to CB merging TFBSs that are located closer than a specified gap parameter, with the default value of this parameter set to 35 bp. To evaluate if the high recall rate of TFBSs unique to CB is due to the merging of close TFBSs, the above analysis was repeated with the gap parameter set to 1 bp. As in the previous results (Figure S3), CB is distinct from the other tools by having high precision and recall for a number of samples clustered at the bottom of the heatmap (Figure S4). However, relative to the findings when the default values were used, using a 1 bp gap parameter results in the maximum recall reducing from 40% to 25%, potentially due to the unmerged matches of CB no longer being unique to the tool. As a result of these findings, unless specified, all analyses performed with CB in this study use the default gap parameter value.

Given the observation that the two clusters of tools in Figure S3 capture complementary sets of functional TFBS in their top-scoring matches, we next explored how these results can be unified into an Ensemble approach. Comparing the global similarity of unique motif matches reveals that the results from FIMO cluster with those of MOODS and MS, while the results from CB are distinct from the other tools (Figure S2 and S3). Due to the similarity of results from FIMO, MOODS, and MS, only the results from FIMO were selected for the Ensemble. FIMO was selected as it achieved the highest precision of the three tools, with the least number of motif matches. All matches found by FIMO were combined with a set of quality matches from CB to overcome the recall problem of FIMO. The top 7000 matches, determined as the optimal number of matches to select based on considerations of precision and recall (Figure S5), were integrated into all matches of FIMO to form the Ensemble set of matches (see “Materials and Methods”).

### Ensemble motif mapping yields additional target genes when characterizing gene regulatory networks from TF perturbation experiments

One of the fields in which motif mapping plays an important role is GRN inference. To validate the applicability of the Ensemble approach to study GRNs in plants, we compared the regulatory links predicted from the motif mapping to lists of genes that are differentially expressed after TF perturbation (DE gene sets). TFs for which perturbation experiments have been conducted covered a wide range of biological processes, such as AGL15 in embryogenesis, AP3 and PI in flower development, BES1 in plant growth and development, FHY3 and PIFs in response to light, WRKY33 in defense response, and EIN3 in response to ethylene (see “Materials and Methods”). To test the recall of the Ensemble, we investigated whether the TFBSs corresponding to the perturbed TF (referred to as the correct TFBSs) were significantly enriched in the promoters of the DE genes (hypergeometric test, FDR corrected p-value≤0.01). Furthermore, the subset of genes from the DE gene set that contained a correct TFBS were compared with experimental ChIP-Seq data for the TF to identify bona fide target genes (see “Materials and Methods”). For 9 of the 10 DE gene sets, a significant enrichment of the correct TFBSs were found for the DE gene sets using the Ensemble. Out of these 9 sets of DE genes, the Ensemble showed better recovery of ChIP-confirmed target genes for 5 sets (PIF4, WRKY33, EIN3, FHY3, and PI) compared to FIMO. For the remaining sets, the rate of recovery was comparable to FIMO. In total, the Ensemble method identified 41 target genes that were missed by FIMO for 10 DE sets (referred to as ‘extra targets’), out of which 32 (78%) were confirmed using ChIP-Seq datasets for the respective TFs (Table S3). For WRKY33 and PI, the Ensemble yielded the largest number of additional ChIP-confirmed target genes (16 for WRKY33 and 8 for PI). Moreover, the FIMO matches lacked a significant enrichment of the WRKY33 motif for WRKY33 perturbed genes. The target genes of WRKY33 which were missed by FIMO included ZFAR1/CZF1 (AT2G40140) and ERF1 (AT3G23240), which are both involved in defense response to biotic stimulus (Table 2). Other examples of target genes detected by Ensemble and missed by FIMO included: HEC1 (AT5G67060) in the PI DE gene set, a well-known TF involved in gynoecium development (Gremski et al., 2007), and BLH1 (AT2G35940) in the FHY3 DE set, a gene known to be involved in the response to far-red light (Staneloni et al., 2009). A detailed intersection of the Ensemble motif matches, the DE genes after TF perturbation, and the ChIP targets is shown in Supplemental Figure S6.

**Table 2.** List of ChIP confirmed targets only identified by the Ensemble approach when comparing enriched TFBSs with perturbed DE gene sets.

| TF | #Extra ChIP confirmed targets | ChIP confirmed targets |
| --- | --- | --- |
| WRKY33 | 16 | AT3G23240(ERF1), AT1G14350(MYB124), AT1G51700(DOF1), AT2G23320(WRKY15), AT2G43140, AT2G01940(IDD15), AT2G40140(SZF2), AT3G55950 (CCR3), AT2G36960(TKI1), AT3G55980(SZF1), AT3G27785(MYB118), AT4G11070(WRKY41), AT4G01250(WRKY22), AT5G56960, AT5G24110(WRKY30), AT5G56550 |
| FHY3 | 6 | AT2G35940(BLH1), AT1G13020(EIF4B2), AT1G35460(bHLH80), AT2G33860(ARF3), AT2G39130, AT5G01780 |
| PI | 8 | AT5G67060(HEC1), AT1G08570(ACHT4), AT1G12240(VI2), AT2G19110(HMA4), AT3G56360, AT3G56370, AT4G01120(bZIP54), AT5G07350(Tudor1) |

### An improved protocol to identify gene regulatory networks starting from accessible chromatin regions

The identification of highly accessible open chromatin regions throughout the genome helps to determine the location of potential regulatory elements. Recent advancements in experimental technologies have allowed researchers to map open chromatin regions in specific plant cell types (Maher et al., 2018; Sijacic et al., 2018). However, identifying which TFs are likely to bind these regions, and how they affect gene expression, is still a major challenge. A recent study used ATAC-Seq to identify transposase hypersensitive sites (THSs) specific to stem cells of the shoot apical meristem (SAM) and leaf mesophyll cells (Sijacic et al., 2018). To identify potential TFs that bind to these THSs, Sijacic and co-workers used *de novo* motif discovery to identify over-represented motifs in these regions. Motifs found *de novo* were compared to known TF motifs to identify potential regulators. The genomic locations of these over-represented motifs, determined using FIMO, were then assigned to the closest gene to identify potential cell type-specific target genes of the associated TFs. By selecting TFs showing cell type-specific expression, measured using high rank ratios (RR) in each cell (see “Materials and Methods”), and that also had at least one *de novo* motif assigned to it, Sijacic et al. reported 23 and 128 TFs in SAM stem and leaf mesophyll cells, respectively.

This traditional pipeline, besides having multiple steps to identify cell type-specific GRNs, is dependent on *de novo* motif discovery tools and parameters. Furthermore, linking motifs found *de novo* with known TFs can be challenging (Castro-Mondragon et al., 2017). In order to overcome some of these problems, we developed a novel protocol in which the enrichment of TFBSs was directly compared against a set of 2,132 SAM stem cell and 1,508 mesophyll-specific THSs, to identify putative regulators and targets (see “Materials and Methods”).

Starting from 59 SAM stem cell-specific and 158 mesophyll cell-specific TFs having high RR, we determined which motifs were significantly enriched in the corresponding cell type THSs using both FIMO and Ensemble motif mappings (see “Materials and Methods”). The Ensemble approach reported a larger number of significantly enriched TFBS compared to FIMO in both cell types. Of 59 TF motifs mapped, 29 were significantly enriched in SAM stem cell-specific THSs when Ensemble mappings were used, whereas 25 motifs were enriched when FIMO motif mappings were used. Whereas 13 motifs correspond to TFs also reported in the original study, 16 of the 29 significantly enriched motifs from the Ensemble set corresponded to TFs that were not described by Sijacic et al. (2018). These TFs include BRC2 from the TCP family, AGL24, AGL27, AGL31, and AGL70 belonging to the MADS family, IDD15 from the C2H2 family and additional TFs from the zf-HD and C2C2-Dof families. For mesophyll cells, 55% (87 out of 158) of the motifs were enriched for mesophyll THS regions using Ensemble motif mappings, whereas for FIMO only 48% (77 out of 158) of the motifs showed a significant enrichment. Eleven of the 87 TFs found enriched using Ensemble motif mappings were not reported by Sijacic et al., and included DREB2, AT1G33760, and RAP2.4 belonging to the AP2-EREBP family, CCA1 and LCL1 from a MYB-related family, AT5G50915 from the bHLH family, bZIP60 from the bZIP family, SPL13 from the SBP family, AT1G14580 from the C2H2 family, and WRKY30 from the WRKY family.

The TFs binding to all enriched motifs identified using the novel protocol in both SAM stem and mesophyll cells were compared with the previously reported 23 and 128 TFs in the respective cell types (Sijacic et al., 2018). Results from the Ensemble method showed enrichment for 13 of 23 motifs, whereas FIMO TFBSs were enriched in 11 of 23 cases. Similarly, for mesophyll cells, 76 and 69 of 128 TF motifs were found to be enriched using the Ensemble and FIMO, respectively (Supplemental Table S4). Overall, our one-step protocol identified 116 regulators showing both significant TFBS enrichment for THSs and cell type-specific expression, of which 23% (n=27) were not reported in the original study. Conversely, for 62 TFs reported by Sijacic and co-workers no significant TFBS enrichment was found using our protocol, suggesting the corresponding motifs do not occur more in the THSs than expected by chance.

To understand how the choice of motif mapping tool affects GRN construction, we investigated the differences between the Ensemble and FIMO motif mappings based on the putative target genes they identify. In total, the Ensemble identified 6,917 targets for 29 significantly enriched motifs in SAM stem cells, whereas FIMO identified 6,428 targets. To determine whether the extra targets identified using the Ensemble are potentially functional, we evaluated their gene expression in each cell type. We initially counted how many of the targets exhibit a two-fold expression difference (|log_2_(RR)| > 1) in either of the cell types. Out of 489 extra targets identified by the Ensemble approach in SAM stem cells, 171 genes (35%) were expressed and 93 genes (19%) showed cell type-specific expression (−log2(RR) > 1; Supplemental Table S4). The fraction of Ensemble-unique target genes which are expressed in a cell type-specific manner are consistent across the different TFs (Figure 4A, TFs labelled blue indicate new regulators). Nine of the cell type-specific genes show more than 6-fold higher expression in SAM stem cells and are therefore likely to be bona fide target genes within the SAM stem cell-specific GRN (Table 3). Three of these nine genes (AT4G11211, AT5G02450, and AT5G13340) lack experimental evidence about their biological function. The remaining 6 genes are known to be involved in a number of processes based on experimental Gene Ontology annotations: primary root development (ATHB13), xylem development (KNAT1), response to cold (DIN10), salt stress and abscisic acid (GASA14), and defense response to bacterium (ERD5, TGA4). These genes are regulated by a diverse array of TFs, such as IDD7, TCP7, ALC, AGL70, DAG2, AGL27, BRC2, JKD, KNAT1, AGL31, and AGL24, that were both described in the original study or identified here. Interestingly, KNAT1 and TGA4, being TF themselves, are regulated by multiple TFs (IDD7 and JKD regulate KNAT1, ALC and KNAT1 regulate TGA4), suggesting some new transcriptional cascades in the SAM stem cell-specific GRN (Figure 4B).

**Table 3.** List of bona fide targets identified in SAM stem and mesophyll cells using the novel TFBS enrichment protocol.

| **Cell type** | **# targets showing cell type-specific expression** | **Target genes** |
| --- | --- | --- |
| SAM stem | 9 | AT1G69780 (ATHB13), AT3G30775 (ERD5), AT4G08150 (KNAT1), AT4G11211, AT5G20250 (DIN10), AT5G13340, AT5G10030 (TGA4), AT5G02450, AT5G14920 (GASA14) |
| Mesophyll | 29 | AT1G14580, AT1G17420 (LOX3), AT1G19450, AT1G51090, AT1G59870 (PDR8), AT1G61890, AT1G72520 (LOX4), AT1G77760 (NR1), AT2G15020, AT2G26530 (AR781), AT2G27310, AT2G36990 (SIG6), AT2G39200 (MLO12), AT3G21670 (NPF6.4), AT3G22060, AT3G24190, AT3G51660, AT3G51895 (AST12), AT3G54020 (AtIPCS1), AT4G02970 (AT7SL-1), AT4G12005, AT4G21570, AT4G34410 (RRTF1), AT5G41740, AT5G42650 (AOS), AT5G44070 (ARA8), AT5G49520 (WRKY48), AT5G49730 (FRO6), AT5G54165 |

**Figure 4.**
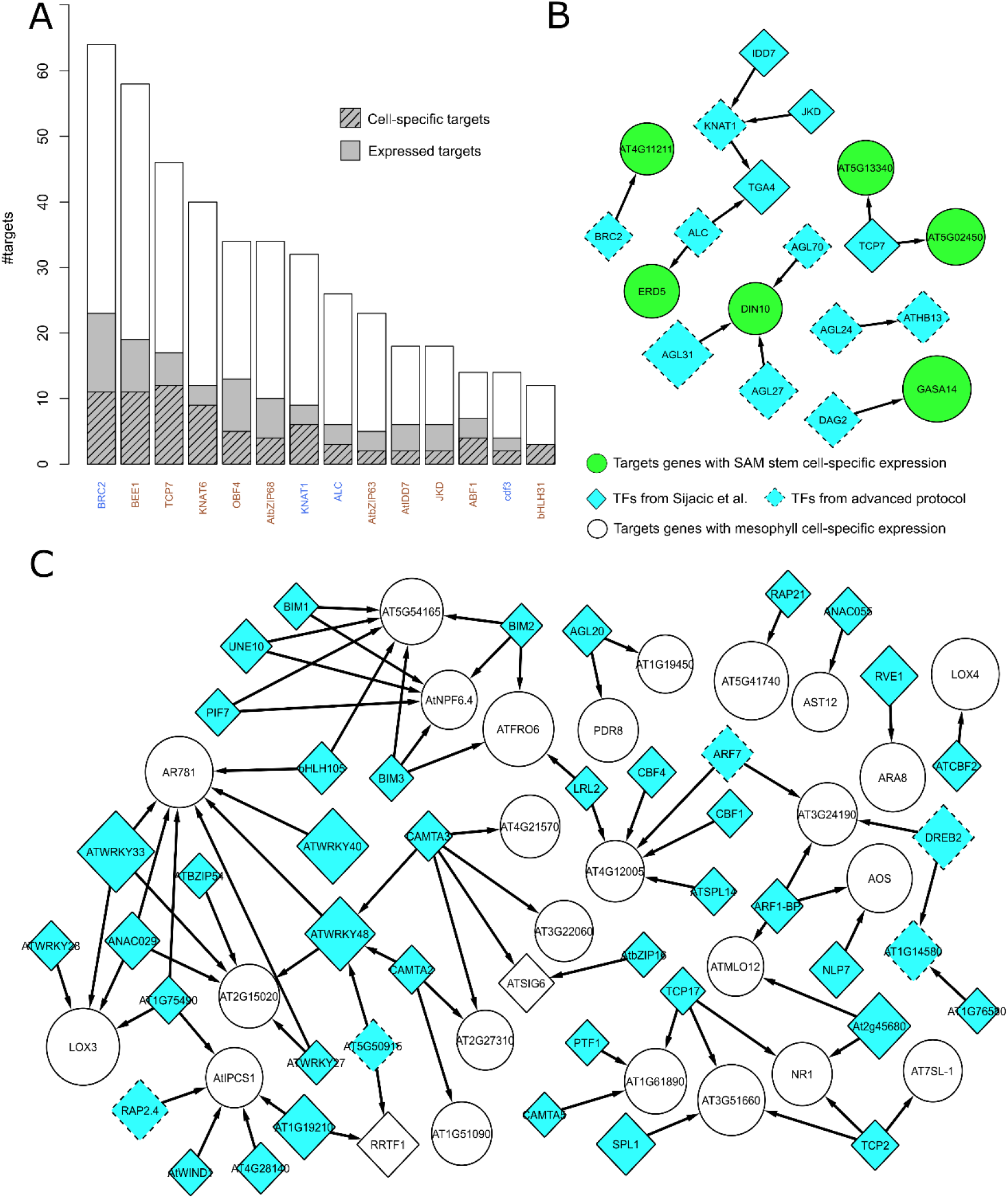
Results of the new protocol to identify potential regulators in SAM stem and mesophyll cell-specific ATAC-seq regions. (A) Barplot showing extra target genes obtained using the Ensemble approach for SAM stem cell. Grey sections show how many of the extra targets have expression in either of the two cell types. Shaded regions show the target genes that are specific to SAM stem cells. TFs labelled in brown are the TFs reported in Sijacic et al., 2018 and TFs labelled in blue are the TFs only identified using the new protocol. Only the TFs that have ≥ 10 extra targets are shown. (B) and (C) AGRN showing SAM stem and mesophyll cell-specific targets identified by enriched TFs respectively. Round green/white refer to the genes that have higher expression in SAM stem/mesophyll cell and diamonds represent TFs. All nodes that have an incoming edge are target genes having high SAM stem/mesophyll cell-specific expression. The size of each node corresponds to the RR of the gene in respective cell type.

For mesophyll cells, 574 of 1,660 new target genes (35%) were expressed in either of the cell types, which is a similar fraction as that reported for SAM stem cells. The percentage of cell type-specific targets identified using the Ensemble motif mappings was 24% for mesophyll cells, corresponding to 402 identified targets with higher expression (−log2(RR) < −1) only in mesophyll cells. 29 of these genes had more than 6-fold expression in mesophyll cells (Table 3). The mesophyll cell-specific GRN of highly expressed genes contained 77 regulatory interactions between 29 targets and 42 TFs, with many of these new target genes being regulated by multiple TFs (Figure 4C). Several of the new target genes have a role in hormone signaling, such as AOS, LOX3, and LOX4, reported to be jasmonic acid responsive, and RRTF1, involved in ethylene biosynthesis. Examples of new unknown target genes are AT5G54165 (regulated by BIM1, BIM2, BIM3, PIF7, UNE10, and bHLH105), AT3G51660 (regulated by TCP2, TCP17, and SPL1), and AT4G12005 (regulated by ARF7, LRL2, SPL14, CBF1, and CBF4). A complete set of interactions between TFs and target genes in SAM stem and mesophyll cells predicted using the Ensemble TFBS enrichment protocol are available as a Cytoscape network session file (Supplemental Data D1).

## DISCUSSION

Recent technological developments have made it possible to profile the chromatin state of particular cell types with high specificity (Maher et al., 2018; Sijacic et al., 2018). This specificity has extended to the level of single cells, allowing cell-to-cell variability in chromatin accessibility to be assessed. However, the impact of these studies is dependent on determining the biological relevance of the accessible regions, particularly if those regions are not located within genes. In addition, as the cost of sequencing decreases and as long read sequencing technologies improve, the number of available genome sequences will increase. While methods to annotate genes are relatively mature, methods to annotate non-coding, regulatory regions are less so. One method of understanding the relevance of accessible chromatin regions, and of annotating potential regulatory sequences, is to map known TF binding preferences onto DNA sequences to identify likely locations bound by TFs. While many tools exist to perform this mapping, each makes certain biological assumptions, and consequently it can be unclear which tool leads to more reliable results in a particular situation.

In order to address this problem, we performed a detailed evaluation of motif mapping tools to determine regulatory relationships in Arabidopsis. Precision and recall were determined for each tool using ChIP-Seq data to assess true positives, revealing that although vastly different numbers of matches were found for each tool, the ability to identify sites that are supported experimentally was similar when similarly-sized subsets of top scoring matches were taken for each tool. FIMO, which is widely used in the plant science community, gave the best precision within its predicted motif matches, but fails to recover some true motif matches due to its stringent settings. Using a benchmark dataset consisting of 40 TFs, we observed that FIMO and CB offer a complementary view on functional TFBSs. We found that when focusing on top7000 matches, despite having a higher false positive rate than FIMO using default settings, CB identified a set of unique motif matches, up to 38% of which were confirmed by ChIP-Seq data. Combining the results of FIMO and CB into an Ensemble set of motif mappings resulted in improved recall relative to FIMO, when motif enrichment of TF perturbation DE gene sets was performed. Overlap of enriched motifs with the ChIP-Seq datasets revealed that, for 5 of the DE gene sets, the Ensemble identified 32 extra functional targets missed by FIMO. Several of these additional TF-target regulatory interactions identified using the Ensemble approach are supported by literature. In independent WRKY33 perturbation experiments (Birkenbihl et al., 2012; Sham et al., 2017), half of the extra targets identified by the Ensemble approach were also found to be DE between wrky33 mutants and wild type Arabidopsis plants (ERF1, FLP / MYB124, DOF1, WRKY15, WRKY30, SZF2, AT2G43140, AT3G55950). In addition to the known role of WRKY33 in defense responses (Birkenbihl et al., 2012), expression of this TF is also associated with more broad stress conditions such as cold, salinity, wounding, and biotic stress (Ma and Bohnert, 2007). Of the 16 extra WRKY33 targets, MYB124, WRKY15, WRKY22, WRKY30, SZF1, and SZF2 have all been found to play roles in a range of stress responses (Sun et al., 2007; Xie et al., 2010; Vanderauwera et al., 2012; Scarpeci et al., 2013; Kloth et al., 2016), supporting the proposed function of WRKY33 as a central stress response factor. The identification of BLH1 as an additional target of FHY3, which integrates responses to far-red light and abscisic acid signaling (Wang and Deng, 2002; Tang et al., 2013), suggests a role for FHY3 during germination and early seedling development, as BLH1 is known to be involved in ABA-mediated signaling pathway acting during early plant development (Kim et al., 2013). The finding of FLOWERING BHLH 1 (FBH1) / bHLH80 as an additional target of FHY3 is also consistent with the role the gene has in light signaling, as FBH1 has been found to control CONSTANS, a key photoperiod gene, and influence the response of the circadian clock to temperature (Ito et al., 2012; Nagel et al., 2014). Finally, the function of PI as a floral homeotic gene in the shoot apical meristem to ensure correct floral organ determination (Goto and Meyerowitz, 1994) is in line with the regulation of HEC1, a bHLH transcription factor which also acts downstream of WUSCHEL to control stem cell proliferation (Schuster et al., 2014). The supporting literature for these interactions strongly suggests that the additional targets identified by the Ensemble motif mappings are functional.

Next, we introduced a novel protocol to learn GRNs from accessible open chromatin regions profiled using cell type-specific ATAC-Seq. Starting from all TFs for which motif information was available, TFBS enrichment was combined with information about cell type-specific expression to infer GRNs. Both traditional and our new protocol inherently depend on the availability of TF motifs, which is a limitation. However, the protocol employed in this study is independent of both *de novo* motif discovery and similarity searches of *de novo* found motifs against motif databases, which is an important step in traditional pipelines to learn regulatory interactions from open chromatin regions (Sullivan et al., 2014; Maher et al., 2018; Sijacic et al., 2018). Moreover, the protocol not only reduces the number of steps to go from cell type-specific THSs to GRNs, but also identifies TFs missed in the previous study by Sijacic et al. (29 and 87 additional significantly enriched TF motifs in SAM stem cells and leaf mesophyll cells, respectively). Conversely, 62 TFs described in the original study were not found to be enriched using our protocol, suggesting there is still room for improvement to learn complete GRNs starting from cell type accessible regions. In addition to finding new TFs, we observed that the performance of the Ensemble approach surpassed that of FIMO when used to map motifs as part of the protocol reported here. Additional enriched TF motifs were identified using the Ensemble, with 4 additional regulators out of 29 total TFs in SAM stem cells and 10 additional TFs out of 87 in mesophyll cells. A striking addition to the set of TF motifs enriched in the SAM stem cell THSs are those of the MADS-box containing genes MAF1/FLM, MAF2, MAF3, and AGL24. All of these genes have been found to influence flowering time, and have positions within a TF network in the SAM that integrates environmental and developmental signals to control flowering (Yu et al., 2002; de Folter et al., 2005; Werner et al., 2005; Rosloski et al., 2010; Capovilla et al., 2017). The motifs of these TFs were found enriched in THSs specific to the SAM stem cells, suggesting that signal integration is occurring in the stem cells at the apex. In addition to these motifs, the motifs corresponding to KNAT1 and AtCSP2 were also enriched. Correspondingly, the expression of both genes has previously been found to be localized to the SAM, with KNAT1 being a homeodomain important for leaf morphogenesis and AtCSP2 involved in the transition to flowering and silique development (Lincoln et al., 1994; Nakaminami et al., 2009). In contrast to the SAM, the additional mesophyll cell-specific enriched motifs contain TFs known to be involved with stress responses, the circadian clock, and growth. DREB2 is involved with controlling drought-responsive genes (Sakuma et al., 2006), while WRKY30 has been found to be important for both biotic and abiotic stress responses (Scarpeci et al., 2013). In addition to stress responses, motifs from TFs involved in the age-related flowering time pathway (SPL13) and the circadian clock (AtCCA1) are enriched (Wang and Tobin, 1998; Xu et al., 2016), consistent with the leaf playing a key role in environmental sensing. Finally, ARF7 is an auxin regulated TF which promotes leaf expansion (Wilmoth et al., 2005).

Taken together, the additional enriched motifs identified in the SAM stem cell and leaf mesophyll specific THSs are consistent with the central role of the SAM in flowering time control, and of the leaf responding to stress elicitors and circadian clock entrainment. This demonstrates that the Ensemble-based approach leads to biologically relevant results that contribute towards a more complete picture of the GRNs active in these two tissues, and that might otherwise be missed when using *de novo* motif based methods. In addition, the new target genes identified by the Ensemble, comprising 93 and 402 target genes for SAM stem and mesophyll cells, respectively, were found to be highly expressed in the corresponding cell types, suggesting that the unique regulators as well as their targets identified by the Ensemble are biologically relevant.

In conclusion, we have shown that an integrative approach, utilizing two complementary motif mapping tools, results in improved power to detect functional TFBS relative to FIMO, the most frequently used tool. This approach facilitates more accurate inference of GRNs and will be especially important as chromatin accessibility data continues to be collected. While motif mapping alone is insufficient to accurately map functional regulatory interactions, determining likely positions can help direct future experimental work. A supplemental website offering the Ensemble TFBS mapping results for 1,793 TF motifs corresponding to 916 Arabidopsis TFs is available at http://bioinformatics.psb.ugent.be/cig_data/motifmappings_ath/as a file in BED format.

## MATERIALS AND METHODS

### Collection of TFBS

The motif collection used for this analysis consisted of 66 Arabidopsis position weight matrices (PWMs) representing 40 TFs from different sources including CisBP (Weirauch et al., 2014), Franco-Zorilla and co-workers (Franco-Zorrilla et al., 2014), Plant Cistrome Database (O’Malley et al., 2016), and JASPAR 2016 (Mathelier et al., 2015). IC of PWMs was calculated using the ‘convert-matrix’ command from rsa-tools version 2012-05-25 with -turn option set to ‘info’ (Turatsinze et al., 2008). TFs were assigned to gene families based on the PlnTFDB 3.0 database (Pérez-Rodríguez et al., 2009).

### PWM mapping using different tools

Four mapping tools that are widely used in the plant science community were evaluated in this study. Cluster-Buster (version ‘Compiled on Sep 22 2017’; (Frith et al., 2003)) was run with the ‘−c’ parameter set to zero, as the other tools do not offer prediction of motif clusters. For FIMO, default parameters were used (meme version 4.11.4; (Grant et al., 2011)). For MOODS (version 1.9.3; (Korhonen et al., 2009)), a p-value threshold of <0.0001 was used to enable comparison to FIMO. This threshold was also used for matrix-scan while all other parameters were set to default (rsa-tools version 2012-05-25; (Turatsinze et al., 2008)). The command lines for the different tools are:

~~~
cbust-linux $PWMfile $seqFile -c 0 -f 1
fimo -o $output $PWMfile $seqFile
moods_dna.py -m $PWMfile -s $seqFile -p $threshold --batch -o $output
matrix-scan -v 1 -matrix_format cb -m $PWMfile -i $seqFile -2str -return limits -return sites -seq_format fasta -o $output
$threshold was set to 0.0001 (default value for FIMO and matrix-scan)
~~~

### Extraction of promoter regions

In addition to the TAIR10 Arabidopsis genome annotation, a set of 5,711 non-coding RNAs (ncRNAs) described by (Liu et al., 2012) were added, resulting in a dataset covering 38,966 genes (Lamesch et al., 2012). For all genes a promoter region 5000bp upstream of the translation start site and 1000bp downstream of the translation end site, including introns, was used. If another gene was present upstream of the gene, the region is cut where this upstream gene starts or ends.

### Estimation of recall, precision, and false positive rate

For each TF, all PWM matches from each mapping tool were overlapped with publicly available TF ChIP-Seq data (Table S5). BEDTools was used to intersect the BED files, using the ‘-f’ option set to 1 for complete overlap (Quinlan, 2014). Precision was calculated as the number of TFBS matches confirmed by ChIP-Seq divided by the total number of matches. Recall was calculated as the number of ChIP-Seq peaks for the studied TF that were covered by motif matches, divided by the total number of ChIP-Seq peaks.

To calculate the false positive rate (FPR) of the motif mappers, shuffled promoters (n=38,966) were generated by shuffling the sequences of the real promoters. The 66 TFBS were mapped to these shuffled promoters. Following (Jayaram et al., 2016), actual negatives (AN) were calculated for every promoter and every motif as the length of the promoter divided by the length of motif. The FPR was then calculated as the number of matches predicted by a specific tool divided by AN. The FPR value for a TFBS is the average over all promoters.

### Selection of optimal number of top scoring matches

To define the set of matches of CB to combine with FIMO, we took progressively larger sets of CB matches and evaluated which set size resulted in the highest F1 score, a metric which combines precision and recall (Figure S5). An optimal F1 score was observed between 7000 and 9000 matches (Figure S5). Based on this observation, the top scoring 7000 matches of were selected to keep an optimal balance between precision and recall for the CB matches. The same number was also used to identify performance of individual mapping tool by considering equal number of top scoring matches for Figure 1.

### Enrichment on DE genes after TF perturbation

10 publicly available DE gene sets after TF perturbation were used to determine motif enrichment (Table S6). We determined, for each TF, the number of DE genes with a proximal TFBS. The significance of this overlap was determined using the hypergeometric distribution. For each enriched motif, the multiple testing corrected p-value (or q-value) of enrichment is determined using the Benjamini-Hochberg correction. Only q-values ≤0.01 were considered significant. For the motifs that are both enriched in the DE and correspond to the perturbed TF, the subset of genes having that motif were retrieved and compared with TF ChIP-Seq binding data (denoted ‘ChIP confirmed hits’ in Table 2). The ChIP-Seq data sets used are the same as discussed in the section, Table S5.

### Case study on cell type-specific transposase hypersensitive sites (THSs)

Based on the ATAC-Seq datasets from (Sijacic et al., 2018), we defined a set of THSs for stem cell and mesophyll cells. Candidate regulators were predicted using the TFBS information present in the mapping file. We identified a set of specific THSs for both cell types, based on a 2-fold (or higher) difference in the ratio between the stem cell and mesophyll counts, yielding two region files with 2,132 stem cell THSs and 1,508 mesophyll THSs (Table S2). Using the TFBS mappings from FIMO and Ensemble, the significance of the overlap between a specific TFBS and a THS region file was assessed. To select the TF motifs for enrichment analysis, RR for each gene was computed by considering expression ranks from Sijacic et al. (2018). RR was calculated as the ratio of expression rank in stem and mesophyll cells. Genes with −log2(RR) > 1 were called SAM stem cell-specific and −log2(RR) < −1 were called mesophyll specific genes. After this selection, 59 and 158 TFs for the SAM stem cell and mesophyll cell respectively were considered for the analysis. These TFs included the TFs reported in Sijacic et al. (2018).

The THS region file and the mapped TFBSs for a given tool (after running BEDTools merge per TFBS) were formatted as BED files and the overlap between both files was determined using the BEDTools function intersectBed with ‘-u’ parameter and the ‘–f’ parameter set to 0.5. As such, we obtained for each THS region file and each TFBS an observed number of mapped TFBSs overlapping with THSs. To determine the significance of this observed overlap, the expected amount of overlapping TFBS with the same THS region file was determined by shuffling the TFBS mapping bed file 1000 times, using shuffleBed with the ‘-noOverlapping’ option enabled across the pre-defined promoter regions (described in “Extraction of promoter regions” section). The overlap with the THS region file was determined for each shuffled file and the median number of TFBS over all shuffled files was used as a measure for the expected overlap. This estimation was used to calculate the fold enrichment, defined as the ratio between observed overlap and expected overlap by chance. An empirical p-value was determined by counting how many times the expected overlap was bigger than or equal to the observed overlap. Only cases where p-value ≤0.01 were considered as significant.

Command line for the pipeline

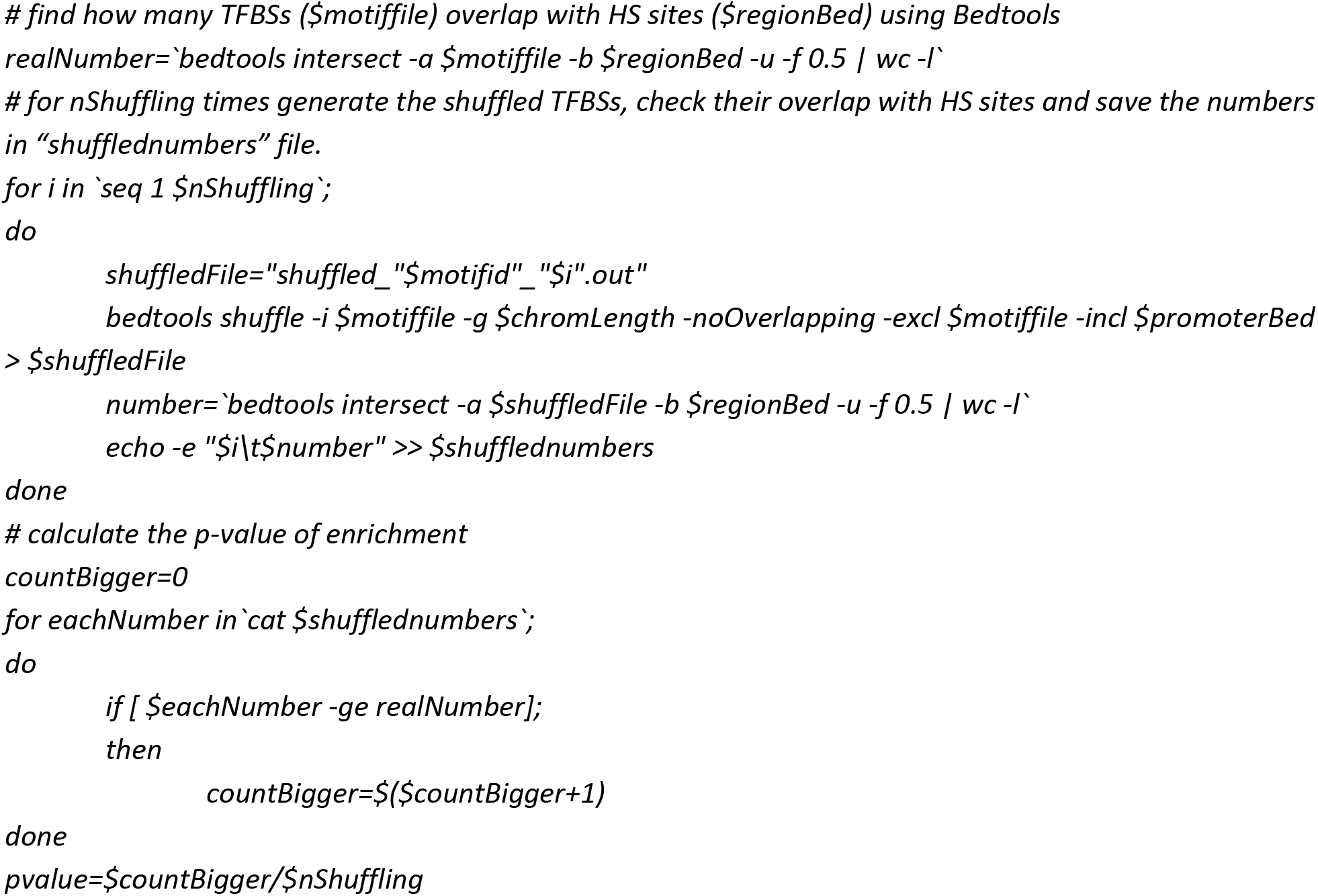

## SUPPLEMENTAL INFORMATION

Supplemental Figure S1. TF level performance of TFBS mapping tools.

Supplemental Figure S2. TF level performance of unique matches considering pairwise combinations of tools for top-scoring 7000 matches.

Supplemental Figure S3. TF level performance of unique matches considering one tool against all other tools for top-scoring 7000 matches.

Supplemental Figure S4. TF level performance of unique matches considering one tool against all other tools for top-scoring 7000 matches and CB gap parameter set to 1.

Supplemental Figure S5. The relationship between F1 score and subset size suggests the top 7000 highest scoring matches of Cluster-Buster should be used in the Ensemble.

Supplemental Figure S6. Overlap analysis for perturbation experiments.

Supplemental Figure S7. Cartoon for an improved protocol to identify GRNs starting from accessible chromatin regions.

Supplemental Table S1. List of publications in plant science community using different mapping tools.

Supplemental Table S2. Overview of 66 TF motifs selected to evaluate the performance of motif mapping tools.

Supplemental Table S3. TFBS enrichment results for DE gene sets.

Supplemental Table S4. List of TFs considered for ATAC-seq case-study with the distribution of their target genes in stem and mesophyll cells

Supplemental Table S5. Overview of TF ChIP-Seq datasets used for estimation of precision and recall

Supplemental Table S6. Overview of DE gene sets after TF perturbation used for the case-study

Supplemental Data D1: Cystoscope session file with GRNs in SAM stem and mesophyll cells described in the case-study.

## ACKNOWLEDGMENTS

We thank Francois Bucchini for the technical assistance to set up the supplemental website.

## REFERENCES

Baxter L, Jironkin A, Hickman R, Moore J, Barrington C, Krusche P, Dyer NP, Buchanan-Wollaston V, Tiskin A, Beynon J, Denby K, Ott S (2012) Conserved noncoding sequences highlight shared components of regulatory networks in dicotyledonous plants. Plant Cell 24: 3949–3965

Birkenbihl RP, Diezel C, Somssich IE (2012) Arabidopsis WRKY33 is a key transcriptional regulator of hormonal and metabolic responses toward Botrytis cinerea infection. Plant Physiol 159: 266–285

Burgess DG, Xu J, Freeling M (2015) Advances in understanding cis regulation of the plant gene with an emphasis on comparative genomics. Curr Opin Plant Biol 27: 141–147

Capovilla G, Symeonidi E, Wu R, Schmid M (2017) Contribution of major FLM isoforms to temperature-dependent flowering in Arabidopsis thaliana. J Exp Bot 68: 5117–5127

Castro-Mondragon JA, Jaeger S, Thieffry D, Thomas-Chollier M, van Helden J (2017) RSAT matrix-clustering: dynamic exploration and redundancy reduction of transcription factor binding motif collections. Nucleic Acids Res 45: e119

de Folter S, Immink RG, Kieffer M, Parenicova L, Henz SR, Weigel D, Busscher M, Kooiker M, Colombo L, Kater MM, Davies B, Angenent GC (2005) Comprehensive interaction map of the Arabidopsis MADS Box transcription factors. Plant Cell 17: 1424–1433

Franco-Zorrilla JM, López-Vidriero I, Carrasco JL, Godoy M, Vera P, Solano R (2014) DNA-binding specificities of plant transcription factors and their potential to define target genes. Proceedings of the National Academy of Sciences 111: 2367–2372

Franco-Zorrilla JM, Solano R (2017) Identification of plant transcription factor target sequences. Biochim Biophys Acta 1860: 21–30

Frith MC, Li MC, Weng Z (2003) Cluster-Buster: Finding dense clusters of motifs in DNA sequences. Nucleic Acids Res 31: 3666–3668

Goto K, Meyerowitz EM (1994) Function and regulation of the Arabidopsis floral homeotic gene PISTILLATA. Genes Dev 8: 1548–1560

Grant CE, Bailey TL, Noble WS (2011) FIMO: scanning for occurrences of a given motif. Bioinformatics 27: 1017–1018

Gremski K, Ditta G, Yanofsky MF (2007) The HECATE genes regulate female reproductive tract development in Arabidopsis thaliana. Development 134: 3593–3601

Haudry A, Platts AE, Vello E, Hoen DR, Leclercq M, Williamson RJ, Forczek E, Joly-Lopez Z, Steffen JG, Hazzouri KM (2013) An atlas of over 90,000 conserved noncoding sequences provides insight into crucifer regulatory regions. Nature genetics 45: 891–898

Heyndrickx KS, Van de Velde J, Wang C, Weigel D, Vandepoele K (2014) A functional and evolutionary perspective on transcription factor binding in Arabidopsis thaliana. Plant Cell 26: 3894–3910

Hickman R, Van Verk MC, Van Dijken AJH, Mendes MP, Vroegop-Vos IA, Caarls L, Steenbergen M, Van der Nagel I, Wesselink GJ, Jironkin A, Talbot A, Rhodes J, De Vries M, Schuurink RC, Denby K, Pieterse CMJ, Van Wees SCM (2017) Architecture and Dynamics of the Jasmonic Acid Gene Regulatory Network. Plant Cell 29: 2086–2105

Ito S, Song YH, Josephson-Day AR, Miller RJ, Breton G, Olmstead RG, Imaizumi T (2012) FLOWERING BHLH transcriptional activators control expression of the photoperiodic flowering regulator CONSTANS in Arabidopsis. Proc Natl Acad Sci U S A 109: 3582–3587

Jayaram N, Usvyat D, AC Rm (2016) Evaluating tools for transcription factor binding site prediction. BMC Bioinformatics

Kim D, Cho YH, Ryu H, Kim Y, Kim TH, Hwang I (2013) BLH1 and KNAT3 modulate ABA responses during germination and early seedling development in Arabidopsis. Plant J 75: 755–766

Kloth KJ, Wiegers GL, Busscher-Lange J, van Haarst JC, Kruijer W, Bouwmeester HJ, Dicke M, Jongsma MA (2016) AtWRKY22 promotes susceptibility to aphids and modulates salicylic acid and jasmonic acid signalling. J Exp Bot 67: 3383–3396

Korhonen J, Martinmaki P, Pizzi C, Rastas P, Ukkonen E (2009) MOODS: fast search for position weight matrix matches in DNA sequences. Bioinformatics 25: 3181–3182

Kulkarni SR, Vaneechoutte D, Van de Velde J, Vandepoele K (2018) TF2Network: predicting transcription factor regulators and gene regulatory networks in Arabidopsis using publicly available binding site information. Nucleic Acids Res 46: e31

Lamesch P, Berardini TZ, Li D, Swarbreck D, Wilks C, Sasidharan R, Muller R, Dreher K, Alexander DL, Garcia-Hernandez M, Karthikeyan AS, Lee CH, Nelson WD, Ploetz L, Singh S, Wensel A, Huala E (2012) The Arabidopsis Information Resource (TAIR): improved gene annotation and new tools. Nucleic Acids Res 40: D1202–1210

Lincoln C, Long J, Yamaguchi J, Serikawa K, Hake S (1994) A knotted1-like homeobox gene in Arabidopsis is expressed in the vegetative meristem and dramatically alters leaf morphology when overexpressed in transgenic plants. Plant Cell 6: 1859–1876

Liu J, Jung C, Xu J, Wang H, Deng S, Bernad L, Arenas-Huertero C, Chua NH (2012) Genome-wide analysis uncovers regulation of long intergenic noncoding RNAs in Arabidopsis. Plant Cell 24: 4333–4345

Lu Z, Hofmeister BT, Vollmers C, DuBois RM, Schmitz RJ (2017) Combining ATAC-seq with nuclei sorting for discovery of cis-regulatory regions in plant genomes. Nucleic Acids Res 45: e41

Ma S, Bohnert HJ (2007) Integration of Arabidopsis thaliana stress-related transcript profiles, promoter structures, and cell-specific expression. Genome Biol 8: R49

Maher KA, Bajic M, Kajala K, Reynoso M, Pauluzzi G, West DA, Zumstein K, Woodhouse M, Bubb K, Dorrity MW, Queitsch C, Bailey-Serres J, Sinha N, Brady SM, Deal RB (2018) Profiling of Accessible Chromatin Regions across Multiple Plant Species and Cell Types Reveals Common Gene Regulatory Principles and New Control Modules. Plant Cell 30: 15–36

Mathelier A, Fornes O, Arenillas DJ, Chen C-y, Denay G, Lee J, Shi W, Shyr C, Tan G, Worsley-Hunt R (2015) JASPAR 2016: a major expansion and update of the open-access database of transcription factor binding profiles. Nucleic acids research: gkv1176

Michael TP, Mockler TC, Breton G, McEntee C, Byer A, Trout JD, Hazen SP, Shen R, Priest HD, Sullivan CM, Givan SA, Yanovsky M, Hong F, Kay SA, Chory J (2008) Network discovery pipeline elucidates conserved time-of-day-specific cis-regulatory modules. PLoS Genet 4: e14

Muino JM, de Bruijn S, Pajoro A, Geuten K, Vingron M, Angenent GC, Kaufmann K (2016) Evolution of DNA-Binding Sites of a Floral Master Regulatory Transcription Factor. Mol Biol Evol 33: 185–200

Nagel DH, Pruneda-Paz JL, Kay SA (2014) FBH1 affects warm temperature responses in the Arabidopsis circadian clock. Proc Natl Acad Sci U S A 111: 14595–14600

Nakaminami K, Hill K, Perry SE, Sentoku N, Long JA, Karlson DT (2009) Arabidopsis cold shock domain proteins: relationships to floral and silique development. J Exp Bot 60: 1047–1062

O’Malley RC, Huang S-sC, Song L, Lewsey MG, Bartlett A, Nery JR, Galli M, Gallavotti A, Ecker JR (2016) Cistrome and Epicistrome Features Shape the Regulatory DNA Landscape. Cell 165: 1280–1292

Pérez-Rodríguez P, Riano-Pachon DM, Corrêa LGG, Rensing SA, Kersten B, Mueller-Roeber B (2009) PlnTFDB: updated content and new features of the plant transcription factor database. Nucleic acids research: gkp805

Quinlan AR (2014) BEDTools: The Swiss-Army Tool for Genome Feature Analysis. Curr Protoc Bioinformatics 47: 11 12 11–34

Rosloski SM, Jali SS, Balasubramanian S, Weigel D, Grbic V (2010) Natural diversity in flowering responses of Arabidopsis thaliana caused by variation in a tandem gene array. Genetics 186: 263–276

Sakuma Y, Maruyama K, Osakabe Y, Qin F, Seki M, Shinozaki K, Yamaguchi-Shinozaki K (2006) Functional analysis of an Arabidopsis transcription factor, DREB2A, involved in drought-responsive gene expression. Plant Cell 18: 1292–1309

Scarpeci TE, Zanor MI, Mueller-Roeber B, Valle EM (2013) Overexpression of AtWRKY30 enhances abiotic stress tolerance during early growth stages in Arabidopsis thaliana. Plant Mol Biol 83: 265–277

Schuster C, Gaillochet C, Medzihradszky A, Busch W, Daum G, Krebs M, Kehle A, Lohmann JU (2014) A regulatory framework for shoot stem cell control integrating metabolic, transcriptional, and phytohormone signals. Dev Cell 28: 438–449

Sham A, Moustafa K, Al-Shamisi S, Alyan S, Iratni R, AbuQamar S (2017) Microarray analysis of Arabidopsis WRKY33 mutants in response to the necrotrophic fungus Botrytis cinerea. PLoS One 12: e0172343

Sijacic P, Bajic M, McKinney EC, Meagher RB, Deal RB (2018) Changes in chromatin accessibility between Arabidopsis stem cells and mesophyll cells illuminate cell type-specific transcription factor networks. Plant J 94: 215–231

Song L, Huang S-sC, Wise A, Castanon R, Nery JR, Chen H, Watanabe M, Thomas J, Bar-Joseph Z, Ecker JR (2016) A transcription factor hierarchy defines an environmental stress response network. Science 354: aag1550

Sparks EE, Drapek C, Gaudinier A, Li S, Ansariola M, Shen N, Hennacy JH, Zhang J, Turco G, Petricka JJ, Foret J, Hartemink AJ, Gordan R, Megraw M, Brady SM, Benfey PN (2016) Establishment of Expression in the SHORTROOT-SCARECROW Transcriptional Cascade through Opposing Activities of Both Activators and Repressors. Dev Cell 39: 585–596

Staneloni RJ, Rodriguez-Batiller MJ, Legisa D, Scarpin MR, Agalou A, Cerdan PD, Meijer AH, Ouwerkerk PB, Casal JJ (2009) Bell-like homeodomain selectively regulates the high-irradiance response of phytochrome A. Proc Natl Acad Sci U S A 106: 13624–13629

Sullivan AM, Arsovski AA, Lempe J, Bubb KL, Weirauch MT, Sabo PJ, Sandstrom R, Thurman RE, Neph S, Reynolds AP (2014) Mapping and dynamics of regulatory DNA and transcription factor networks in A. thaliana. Cell reports 8: 2015–2030

Sun J, Jiang H, Xu Y, Li H, Wu X, Xie Q, Li C (2007) The CCCH-type zinc finger proteins AtSZF1 and AtSZF2 regulate salt stress responses in Arabidopsis. Plant Cell Physiol 48: 1148–1158

Tang W, Ji Q, Huang Y, Jiang Z, Bao M, Wang H, Lin R (2013) FAR-RED ELONGATED HYPOCOTYL3 and FAR-RED IMPAIRED RESPONSE1 transcription factors integrate light and abscisic acid signaling in Arabidopsis. Plant Physiol 163: 857–866

Turatsinze JV, Thomas-Chollier M, Defrance M, van Helden J (2008) Using RSAT to scan genome sequences for transcription factor binding sites and cis-regulatory modules. Nat Protoc 3: 1578–1588

Van de Velde J, Heyndrickx KS, Vandepoele K (2014) Inference of transcriptional networks in Arabidopsis through conserved noncoding sequence analysis. The Plant Cell 26: 2729–2745

Vandepoele K, Casneuf T, Van de Peer Y (2006) Identification of novel regulatory modules in dicotyledonous plants using expression data and comparative genomics. Genome Biol 7: R103

Vandepoele K, Quimbaya M, Casneuf T, De Veylder L, Van de Peer Y (2009) Unraveling transcriptional control in Arabidopsis using cis-regulatory elements and coexpression networks. Plant Physiol 150: 535–546

Vanderauwera S, Vandenbroucke K, Inze A, Van de Cotte B, Muhlenbock P, De Rycke R, Naouar N, van Gaever T, van Montagu MC, Van Breusegem F (2012) AtWRKY15 perturbation abolishes the mitochondrial stress response that steers osmotic stress tolerance in Arabidopsis. Proc Natl Acad Sci U S A 109: 20113–20118

Varala K, Marshall-Colon A, Cirrone J, Brooks MD, Pasquino AV, Leran S, Mittal S, Rock TM, Edwards MB, Kim GJ, Ruffel S, McCombie WR, Shasha D, Coruzzi GM (2018) Temporal transcriptional logic of dynamic regulatory networks underlying nitrogen signaling and use in plants. Proc Natl Acad Sci U S A 115: 6494–6499

Wang H, Deng XW (2002) Arabidopsis FHY3 defines a key phytochrome A signaling component directly interacting with its homologous partner FAR1. EMBO J 21: 1339–1349

Wang ZY, Tobin EM (1998) Constitutive expression of the CIRCADIAN CLOCK ASSOCIATED 1 (CCA1) gene disrupts circadian rhythms and suppresses its own expression. Cell 93: 1207–1217

Weirauch MT, Yang A, Albu M, Cote AG, Montenegro-Montero A, Drewe P, Najafabadi HS, Lambert SA, Mann I, Cook K, Zheng H, Goity A, van Bakel H, Lozano JC, Galli M, Lewsey MG, Huang E, Mukherjee T, Chen X, Reece-Hoyes JS, Govindarajan S, Shaulsky G, Walhout AJM, Bouget FY, Ratsch G, Larrondo LF, Ecker JR, Hughes TR (2014) Determination and inference of eukaryotic transcription factor sequence specificity. Cell 158: 1431–1443

Werner JD, Borevitz JO, Warthmann N, Trainer GT, Ecker JR, Chory J, Weigel D (2005) Quantitative trait locus mapping and DNA array hybridization identify an FLM deletion as a cause for natural flowering-time variation. Proc Natl Acad Sci U S A 102: 2460–2465

Wilmoth JC, Wang S, Tiwari SB, Joshi AD, Hagen G, Guilfoyle TJ, Alonso JM, Ecker JR, Reed JW (2005) NPH4/ARF7 and ARF19 promote leaf expansion and auxin-induced lateral root formation. Plant J 43: 118–130

Xie Z, Li D, Wang L, Sack FD, Grotewold E (2010) Role of the stomatal development regulators FLP/MYB88 in abiotic stress responses. Plant J 64: 731–739

Xu M, Hu T, Zhao J, Park MY, Earley KW, Wu G, Yang L, Poethig RS (2016) Developmental Functions of miR156-Regulated SQUAMOSA PROMOTER BINDING PROTEIN-LIKE (SPL) Genes in Arabidopsis thaliana. PLoS Genet 12: e1006263

Yu CP, Chen SC, Chang YM, Liu WY, Lin HH, Lin JJ, Chen HJ, Lu YJ, Wu YH, Lu MY, Lu CH, Shih AC, Ku MS, Shiu SH, Wu SH, Li WH (2015) Transcriptome dynamics of developing maize leaves and genomewide prediction of cis elements and their cognate transcription factors. Proc Natl Acad Sci U S A112: E2477–2486

Yu H, Xu Y, Tan EL, Kumar PP (2002) AGAMOUS-LIKE 24, a dosage-dependent mediator of the flowering signals. Proc Natl Acad Sci U S A 99: 16336–16341

Zhang W, Zhang T, Wu Y, Jiang J (2012) Genome-wide identification of regulatory DNA elements and protein-binding footprints using signatures of open chromatin in Arabidopsis. The Plant Cell 24: 2719–2731

